# Discovery and Functional Characterization of Salinity-Responsive Promoters in *Pseudomonas putida* KT2440

**DOI:** 10.1101/2025.11.11.687801

**Authors:** Camila Marques de Simone, Guilherme Marcelino Viana de Siqueira, María-Eugenia Guazzaroni

## Abstract

Microbial bioprocesses are associated with environmental advantages, but their freshwater demand remains a challenge. Seawater is abundant and a sustainable alternative; however, high salinity often limits microbial growth and survival. This study aimed to identify biological parts involved in microbial salt stress responses for applications in microbial engineering. Regulatory regions adjacent to genes related to salinity and other industrially relevant conditions in *Pseudomonas putida* KT2440 were cloned into a reporter plasmid for functional evaluation. We identified and characterized the promoters of PP_1833 and PP_4707 as salt-inducible in KT2440, showing fluorescence levels 1.9- and 2.8-fold greater than a standard strong constitutive promoter (P_*j100*_) under 3.5% NaCl (w/v), respectively, while no activity was detected in *Escherichia coli* DH10B. These results provide a basis for developing salt-responsive regulatory circuits in *P. putida* KT2440.

**For table of contents only:** 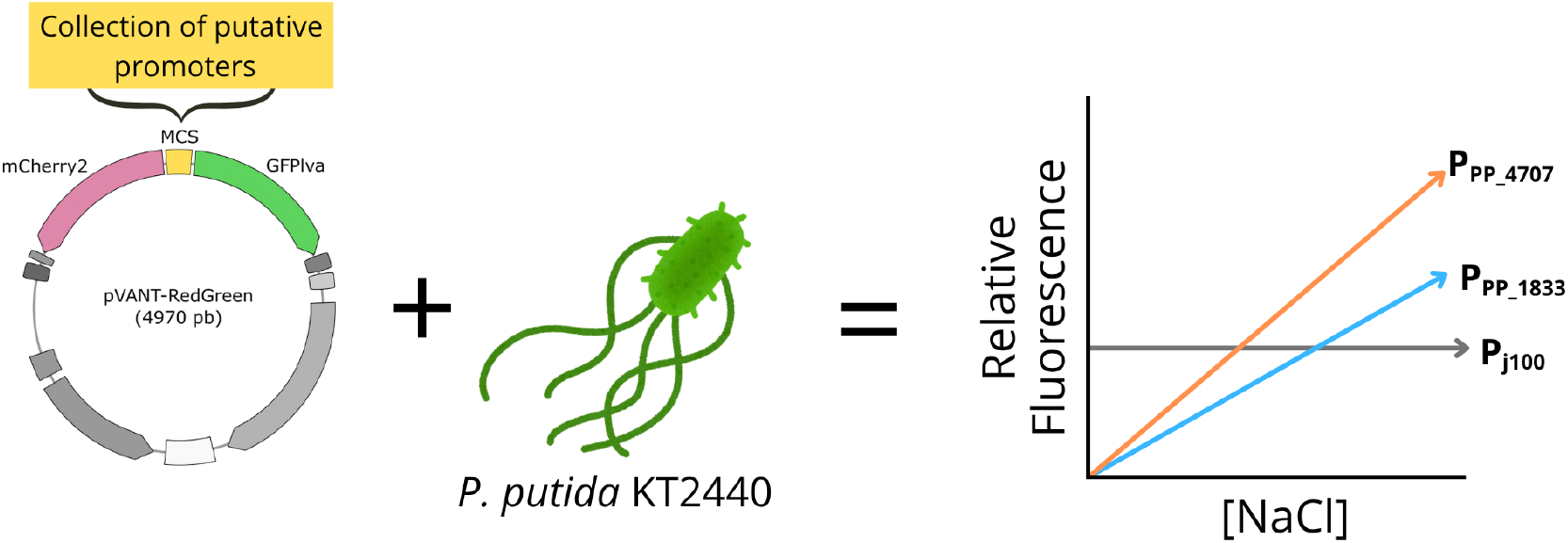

## Introduction

Synthetic biology and industrial biotechnology are key drivers to establishing processes based on the sustainable use of biological resources, upholding the principles of a green and circular bioeconomy. Processes such as microbial fermentations, which are capable of converting waste into value-added products ^1–4^, enable sustainable production routes for a variety of chemicals, including pharmaceuticals, biofuels, plastics, and dyes. However, many bioprocesses still present a high water footprint due to the intensive reliance on freshwater. In biomass biorefining and ethanol production, for instance, water can be consumed at a rate of as much as 9.8 L per liter of product^5^, and this might undermine the benefits of biobased production routes, emphasizing the urgent need for innovative strategies that reduce freshwater dependence while maintaining process efficiency^6,7^.

Seawater represents 97% of the water available in the world^8,9^, and is considered a promising alternative in the face of increasing freshwater scarcity and security concerns. The use of seawater in industrial culture media formulations is not only advantageous to reduce freshwater dependency, but its high salt content also lowers contamination risks while incorporating essential minerals into the fermentation media. As such, different applications for high-salinity bioprocesses have already been reported, such as bioethanol production with marine yeast^10^, cellulosic ethanol with *Saccharomyces cerevisiae, Pichia stipitis*, and *Zymomonas mobilis*^11^, succinic acid by *Actinobacillus succinogenes*^12^, lipids by *Yarrowia lipolytica*^13^ and even baking yeast^14^.

Despite the immense potential of seawater-based microbial processes, such conditions may compromise microbial survival by hindering intracellular water uptake, reducing enzymatic activity, and imposing other osmotic challenges that require specific adaptation mechanisms^10,15^. The salinity of saltwater varies according to location, ranging from less than 3% (w/v) to more than 3.8% (w/v) in certain regions^16^. In standardized synthetic formulations, this corresponds to approximately 35 g/L of salts, predominantly NaCl^12,17,18^. While several halophilic and halotolerant taxa have developed multiple adaptive strategies to overcome these challenges, including modifications in proteins and nucleic acids, accumulation of compatible solutes, and maintenance of stable cell membranes^19,20^, these organisms are often not ideal fermentation hosts since difficulties in their isolation, cultivation, and genetic manipulation limit their scope of application^21^.

Still, several of the mechanisms through which microorganisms respond to environmental stimuli, including transducers, regulatory proteins, and small RNAs, are known and have inspired the engineering of synthetic gene circuits that combine regulatory elements for applications such as metabolite production, bioremediation, and biosensing^22,23^. The success and applicability of these circuits in industrial applications, however, depend not only on genetic design but also on the choice of host. In this context, non-halophilic species such as *Pseudomonas putida* gain relevance by combining ease of manipulation with broad metabolic capacity for producing compounds of interest^24–30^. *P. putida* stands out for its versatile metabolism, lack of virulence factors, and high tolerance to physicochemical stresses^31,32^. The KT2440 strain, in particular, is considered an ideal chassis in synthetic biology due to its robustness and flexibility^33–35^ and well-established genetic manipulation tools^36–41^. Despite being sensitive to elevated concentrations of salt, this species exhibits several osmotic stress response mechanisms that enable growth in such conditions: more than 2,000 genes are differentially expressed under NaCl stress, including permeases, efflux pumps, and transporters regulated by different sigma factors and transcriptional regulators^42^.

Considering this, the present work aimed to identify and characterize promoters related to salt stress in *P. putida* KT2440, with the intention of expanding the toolbox of biological parts to enable the development of genetic circuits for high-salinity cultivation systems in this host.

## Results and Discussion

### *In silico* identification of salt-inducible promoters

The identification of putative promoters associated with genes overexpressed under NaCl stress in this work was based on data from a collection of previously published expression datasets for *P. putida* KT2440 grown under various conditions^43^. We queried this dataset for genes commonly upregulated under salt stress and other stresses relevant to bioprocesses, such as starvation, oxidative stress, and the presence of organic compounds (**Methods**), and investigated the regions contained between their start codon and their nearest ORF, which we refer to as *intergenic regions* (IGR), in search of putative stress-responsive biological parts.

In total, our initial subset of genes included over 500 genes that were upregulated under at least two stress conditions, with only two genes shared across five of them (**Figure 1A**). As we increased the number of intersections, most of the shared ORFs were annotated as hypothetical proteins or proteins containing domains of unknown function (DUFs), which suggested their possible roles as global stress tolerance mechanisms. The two ORFs found to be upregulated in five conditions (PP_1833 and PP_4707) were commonly overexpressed in the NaCl dataset, and were selected as candidates for our screening. These genes encode, respectively, a DUF-3509 domain-containing protein and a BON domain-containing protein, and are annotated in the antisense direction relative to the genomic 5’ → ‘3 orientation (**Figure 1B, Figure 1C**). Interestingly, PP_1833 is neighbored by a gene in the opposite orientation, suggesting the presence of one or two promoters in its IGR (**Figure 1C**).

**Figure 1.**
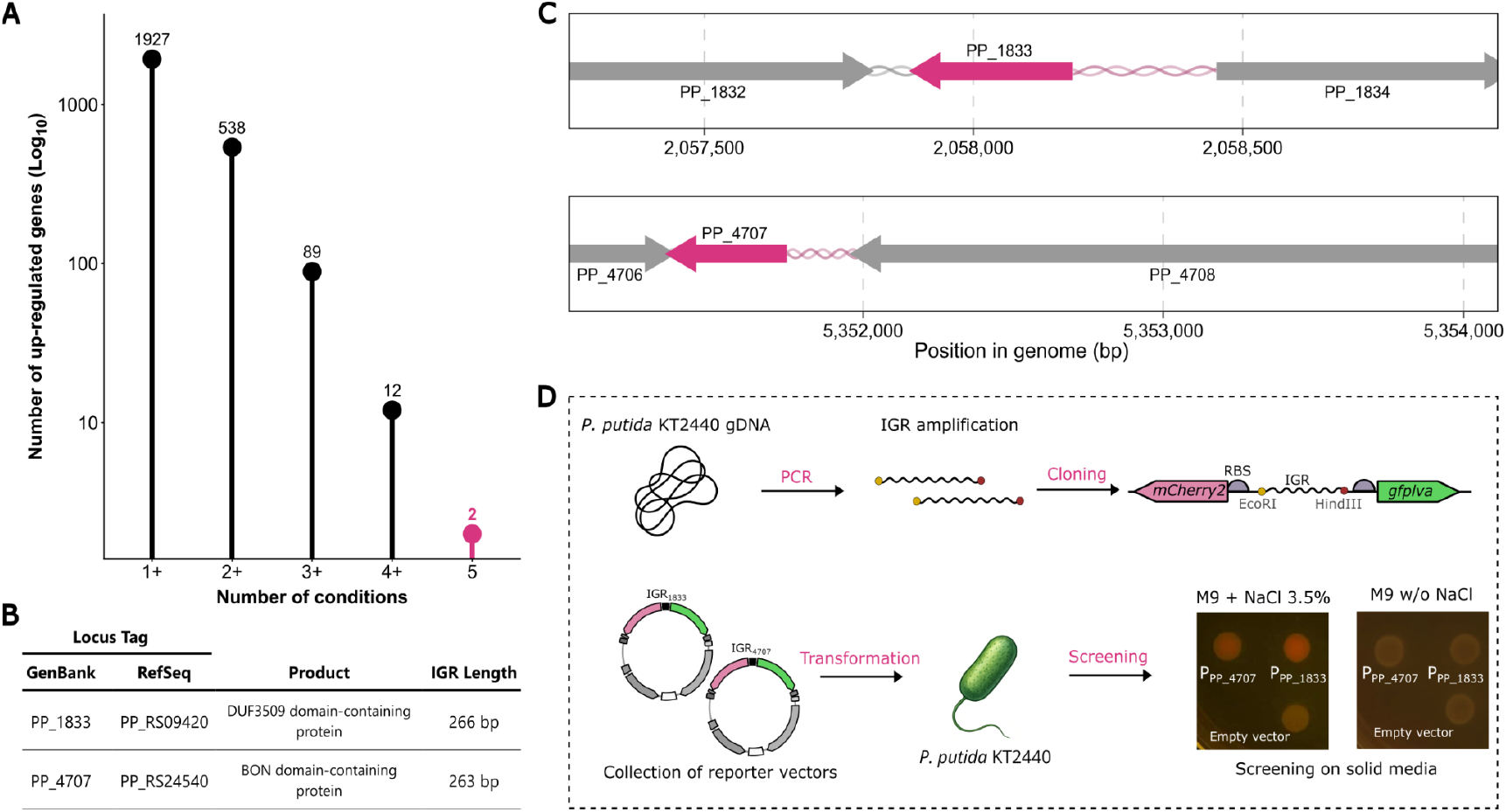
Selection and functional analysis of salt-responsive intergenic regions. (A) Number of upregulated genes detected under one or more stress conditions. Only two genes were up-regulated in all five conditions tested (pink). (B) Summary of genomic features and annotation of PP_1833 and PP_4707. (C) Genomic context of the selected genes, represented in pink with neighboring genes in gray and genomic coordinates in base pairs. (D) Schematic representation of the experimental design used to assess the promoter activity of the intergenic regions (IGRs) of PP_1833 and PP_4707. Each IGR was cloned between the *mCherry2* and *gfplva* reporter genes, and the resulting constructs were evaluated in *P. putida* KT2440. gDNA: genomic DNA

To test whether their IGRs contained suitable stress-responsive promoters, we cloned their intergenic regions into a pVANT-based reporter vector, flanked by reporters *mCherry2* and *gfplva* **(Figure 1D)**, and evaluated their behavior in *P. putida* using solid media (**Methods**). In this first screening, the expected putative native promoters P_PP_1833_ and P_PP_4707_ showed notable induction of the fluorescent reporter in media containing NaCl, with minimal leaky expression when uninduced (**Figure 1D**), indicating they possess desirable traits as salt-inducible promoters, and were therefore subjected to further characterization. While these observed results coincided with expression data, we were not able to observe the dual promoter activity for P_PP_1833_ IGR, suggesting that the regulation of PP_1834 is not affected by the presence of NaCl in the medium under the tested conditions.

### Functional Validation Reveals Salt-Responsive Promoters in P. putida KT2440

To better characterize the profiles of putative P_PP_1833_ and P_PP_4707_ as salt-responsive promoters, pVANT-RG vectors containing IGR sequences were cloned into both *P. putida* KT2440 and *Escherichia coli* DH10B, a well-established model organism and synthetic biology workhorse. Medium turbidity (OD_600_) and fluorescence emission were monitored for strains harboring the reporter vectors in microplate incubations in media containing 0%, 1.7%, and 3.5% NaCl (w/v)—corresponding to a total salt concentration (TS) of 1% (w/v), 2.7% (w/v), and 4.5% (w/v), respectively.

Our results showed two different profiles for *E. coli* and *P. putida* cells with vectors harboring these parts. While all strains showed similar performance when grown under saline conditions (**Supplementary Figure 1**), we observed fluorescence to increase proportionally with salt concentration only when *P. putida* KT2440 was used as the host **(Figure 2**), suggesting that the intergenic regions contain host-specific salt-inducible promoters dependent on regulatory proteins which may not be present or active in *E. coli* under the tested conditions (**Supplementary Figure 2**). For *P. putida*, fluorescence levels in the condition with 3.5% NaCl (w/v) (TS: 4.5% w/v) exceeded those of the positive control (standard strong constitutive promoter *P*_*j100*_), reaching 1.9-fold and 2.8-fold higher values for the P_PP_1833_ and P_PP_4707_ constructs, respectively. Moreover, we were only able to detect expression of *mCherry2*, suggesting that these promoters are unidirectional and active only in the antisense direction of the genome annotation under the tested conditions.

**Figure 2.**
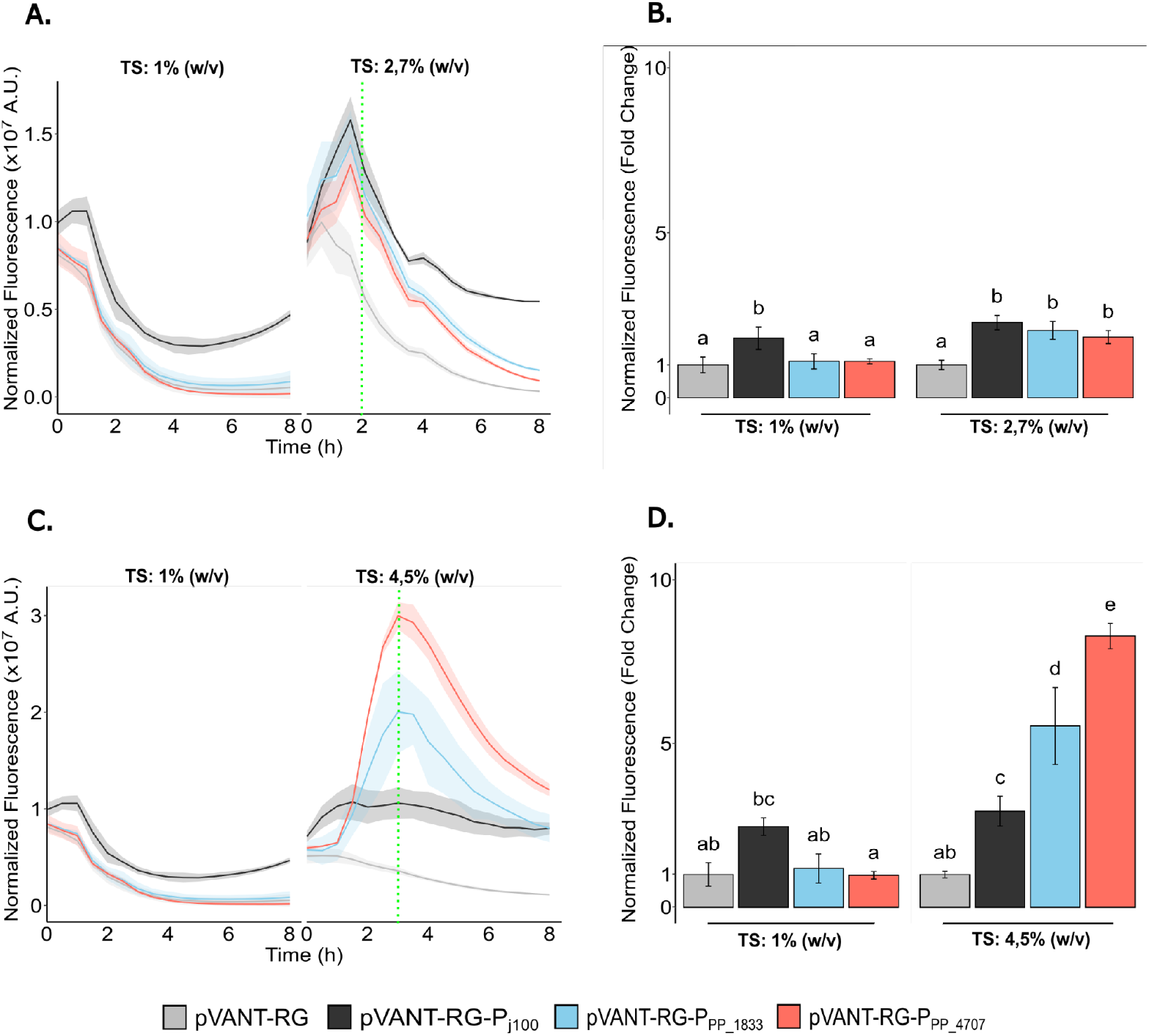
Variation of normalized fluorescence in *P. putida* KT2440. Fluorescence was monitored over 8 hours in microplate assays under different NaCl concentrations (A and C). Specific time points (green line) were analyzed at 2 hours (B) and 3 hours (D), based on the fluorescence behavior observed under saline conditions. Two-way ANOVA followed by Tukey’s post-hoc test revealed statistically significant differences between the constructs containing the P_PP_1833_ and P_PP_4707_ and the plasmid lacking promoters (pVANT-RG) when the strains were cultivated under saline conditions. Comparisons were also made with a positive control: mCherry2 under control of a strong promoter, P_j100_, fused to the fluorescent protein (pVANT-RG-P_j100_). Different significance groups are indicated above the bars. TS: 1% (w/v) = M9; TS: 2.7% (w/v) = M9 + 2.7% NaCl; TS: 4.5% (w/v) = M9 + 3.5% NaCl. A.U. = arbitrary units.

Subsequently, we performed *in silico* prediction of these putative promoters using the software BPROM^44^ and SAPPHIRE^45,46^, identifying specific regions essential for transcription and translation processes. The analysis revealed conserved promoter motifs, including defined −35 and −10 elements, Shine-Dalgarno sequences, and transcriptional and translational start sites, confirming the presence of potential functional promoter architectures within the intergenic regions **(Supplementary Figure 3**).

The contrasting fluorescence patterns observed between *P. putida* KT2440 and *E. coli* DH10B highlight species-specific differences in promoter responsiveness to salinity. Even though both bacteria utilized in this study are Gammaproteobacteria, their specific metabolic machineries are fine-tuned to different environmental conditions and physiological needs. Due to these differences, it is unsurprising that the promoters identified in this study are not readily cross-compatible between them. Several studies have shown that genetic parts often behave differently across bacterial species due to host-specific and context-dependent factors. For instance, promoters and terminators exhibit variable activity between *E. coli* and *P. putida*, with circuit performances also influenced by codon usage, regulatory context, and plasmid backbone^37,47–50^. In the case of the biological parts investigated in this work, both PP_1833 and PP_4707 were recently classified into the “osmotic stress-2” independently modulated (iModulon) group in a study^51^ based on the machine learning-based investigation of *P. putida* KT2440 transcriptome. However, further phenotypic information on PP_1833 and PP_4707 is limited to a few examples indirectly associating them with stress responses^43,52–54^ with their native roles and regulation mechanisms remaining largely poorly understood. Further characterization of these aspects is therefore needed before applying these parts in other bacterial hosts of interest.

While susceptible to stress, *Pseudomonas spp*. have been shown to possess several mechanisms to respond to elevated concentrations of salt^42,52,55^, and some *Pseudomonas* strains can even promote plant growth under saline conditions by mitigating salt stress in their hosts^53,54^, underscoring the broad appeal of investigating the molecular response to salinity in this genus. Our results are consistent with these findings, and In our experiments, *P. putida* KT2440 exhibited moderate yet acceptable growth at 2.7% and even at 4.5% (w/v) total salts concentration (**Supplementary Figure 1**). All this considered, the identification of salt-inducible promoters in *P. putida* KT2440 that we have carried out in this work opens new avenues for the rational design of microbial systems capable of thriving under saline conditions and processes leveraging the use of seawater for media formulation or as an induction system. Future studies should focus on expanding the library of regulatory parts responsive to salinity, including promoters, terminators, ribosome-binding sites, and non-coding RNAs involved in osmotic stress responses, and integrating these elements into synthetic gene circuits that would enable dynamic and tunable control of metabolic pathways during cultivation in seawater-based media, enhancing microbial tolerance while reducing reliance on freshwater in industrial bioprocesses.

## Materials and methods

### Bacterial strains and culture media

In this study, the bacterial strains *E. coli* DH10B, *P. putida* KT2440 were employed. These strains were routinely cultured in M9 minimal medium, composed of 1× M9 salts (42.3 mM Na_2_HPO_4_, 22.0 mM KH_2_PO_4_, 18.7 mM NH_4_Cl and 8.6 mM NaCl), 2 mM MgSO_4_·7H_2_O, 0.1 mM CaCl_2_, 0.1% (w/v) casamino acids and 1% carbon source. Because the M9 medium contains approximately 1% salts, and different concentrations of NaCl were added in the experiments, we adopted the abbreviation “TS” to indicate the total salt content present in the medium. For each experiment, the carbon source was adjusted according to the bacterial strain: *E. coli* DH10B was cultured in M9 medium supplemented with 1% (v/v) glycerol, while *P. putida* KT2440 was grown with 1% (w/v) sodium citrate. All media contained 50 µg·mL^−1^ kanamycin to maintain plasmid selection. Cultures were incubated at 30 °C according to the experimental design. Liquid cultures were grown in an orbital shaker (New Brunswick I26R Refrigerated Incubator Shaker Series, orbital diameter 0.0254 m) at 30 °C, operating at 200 rpm.

### Identification and cloning of salt stress–inducible promoters in P. putida

To identify genes associated with salt stress and their promoters, we utilized the RefSeq-based transcriptome dataset from a previous analysis *P putida* KT2440 expression data built with the reference annotation accession NC_002947.4^43^. The inclusion criteria for conditions in the present work comprised a variety of industrially relevant stressors such as NaCl,H_2_O_2_, Zinc, toluene, ferulic acid and ionic liquids^42,56– 59^, and nutrient limitation^60^. For the purposes of this analysis, a gene was considered to be differentially expressed if its log_2_ fold change value was greater than 1, and the adjusted p-value was lower than 0.05.

After targets were selected, the intergenic regions (IGRs) encompassed between their start codon and the extremity of the nearest annotated ORF were amplified by PCR (**Supplementary Table 1, Supplementary Table 2**) and cloned between the EcoRI and HindIII restriction sites in the multiple cloning site (MCS) of a reporter pVANT vector^37^. The version of the vector used in this work possesses a MCS flanked by *gfplva*^61^ inserted between HindIII and SpeI sites of the pVANT backbone, and *mCherry2*, implemented as the *mCherry2*-L variant^62^, inserted between the PacI and EcoRI restriction sites in the plasmid backbone in the antisense direction in relation to *gfplva*. This pVANT variant is referred to as pVANT-RedGreen or pVANT-RG throughout this manuscript.

For plasmid maintenance and replication, the constructs were first introduced into *E. coli* DH10B. A total of 50 μL of electrocompetent cells were electroporated with 50–100 ng of plasmid DNA using a MicroPulser Electroporator (Bio-Rad, Hercules, CA, USA) under the Ec1 program (1.8 kV) with 0.1 mm cuvettes. Following electroporation, 950 μL of LB medium were added, and the cells were incubated for 1 h at 30 °C to allow recovery before plating on LB agar supplemented with kanamycin (50 µg·mL^−1^). The transformed cells were stored at −80 °C in 20% (v/v) glycerol. For subsequent characterization, the plasmids were introduced into *P. putida* KT2440. Electrocompetent *P. putida* cells were prepared by cultivating the strain in 10 mL of LB medium for 16 h at 30 °C. Cells were harvested by centrifugation at 16,330 × g and 4 °C for 10 min (Eppendorf 5810 R, rotor FA-45-6-30), washed three times with chilled Milli-Q water, and resuspended in approximately 300 μL of chilled Milli-Q water. Aliquots of 50 μL were used immediately for electroporation with 50–100 ng of plasmid DNA, using the same parameters as those applied for *E. coli* DH10B.

An initial test was conducted to qualitatively compare fluorescence activity in *P. putida* KT2440. For this purpose, assays were performed using M9 culture medium supplemented with 1% (w/v) citrate for *P. putida* as carbon source. Transformed strains were first inoculated into 50 mL conical tubes containing 5 mL of M9 liquid medium and incubated in an orbital shaker (New Brunswick I26R Refrigerated Incubator Shaker Series; orbital amplitude: 0.0254 m) at 200 rpm and 30 °C for 16 hours. The optical density at 600 nm (OD_600_) was then measured and adjusted to 1.0 to standardize the cell concentration across all bacterial samples. Subsequently, the cells were diluted in 1× PBS buffer (10 mM sodium phosphate, 137 mM sodium chloride, 2.7 mM potassium chloride; pH 7.4), and 10 μL droplets were spotted onto plates containing M9 solid medium, with or without NaCl supplementation. The salt concentration in the M9 medium was adjusted to 1%, and an additional condition was tested by supplementing the medium with 3.5% (w/v) NaCl (ST: 4.5% (w/v)). After complete drying, the plates were incubated at 30 °C for 16 hours and visually inspected for fluorescence using a Safe Imager™ 2.0 Blue-Light Transilluminator (Invitrogen, Waltham, MA, USA).

### Quantitative Fluorescence assays

Promoter strength was quantified based on *mCherry2* fluorescence measured by spectrophotometry. To this end, the fluorescence of the constructs was evaluated in *E. coli* DH10B and *P. putida* KT2440, testing in both species allowed the evaluation of inherent differences in gene expression. The pVANT-RG construct containing the strong promoter P_j100_^61^ was used as a positive control, whereas the empty vector (lacking promoter insertion) served as a negative control. To assess promoter activity under osmotic stress, *mCherry2* fluorescence was also measured using a microplate reader in M9 medium supplemented with NaCl at different concentrations: 0%, 1.7%, 3.5%, and 5.3% (w/v) — corresponding to total salinity (TS) levels of 1%, 2.7%, 4.5%, and 6.5% (w/v), respectively.

To validate the activity of potential promoters, fluorescence from mCherry2 was measured using a microplate reader in M9 medium supplemented with NaCl at different concentrations (0%, 1.7%, 3.5%, and 5.3% (w/v) — TS: 1%, 2.7%, 4.5%, 6.5% (w/v)). For all experiments, isolated colonies of transformed strains were inoculated into 5 mL of M9 medium with a carbon source (1% (v/v) glycerol for *E. coli* DH10B or 1% (w/v) citrate for *P. putida* KT2440) in 50 mL conical tubes, incubated for 16 h at 30 °C, 200 RPM (orbital amplitude 0.0254 m). Cultures were centrifuged, washed three times with 5 mL of 1X PBS (3,240 × g, 23 °C, 10 min), and resuspended in 5 mL of the same buffer. Aliquots were then transferred to media with different salt concentrations to start with an optical density of 0.05 at 600 nm (OD600) in 200 μL per well.

Samples were dispensed in 96-well opaque microplates with clear bottoms and incubated at 30 °C in a Victor X3 microplate reader (Perkin Elmer). Over eight hours, the reader periodically measured optical density (OD_600_) and fluorescence (Excitation: 587 nm / Emission: 615 nm). Each microplate experiment was conducted with at least three biological replicates, and within each replicate, three technical replicates per sample. Data analysis and statistics were performed using R (version 4.5.1).

## Acknowledgments

The authors would like to thank the lab technicians Thalita Riul Prado and M.Sc Lucas Silva Brito, as well as the MetaGenLab team for their support during the investigation.

## Author Contributions

Conceptualization: M.E.G and G.M.V.S. Data acquisition, processing, and interpretation: C.M.S. and G.M.V.S. Preparation of figures: C.M.S. and G.M.V.S. Initial draft of manuscript: C.M.S. Writing, review, editing of manuscript: C.M.S., G.M.V.S. and M.E.G. Raised funds: M.E.G. All authors have read, given feedback, and approved the manuscript for publication.

