## Supplementary material for "Discovery and Functional Characterization of Salinity-Responsive Promoters in *Pseudomonas putida* KT2440"

### Supporting Information

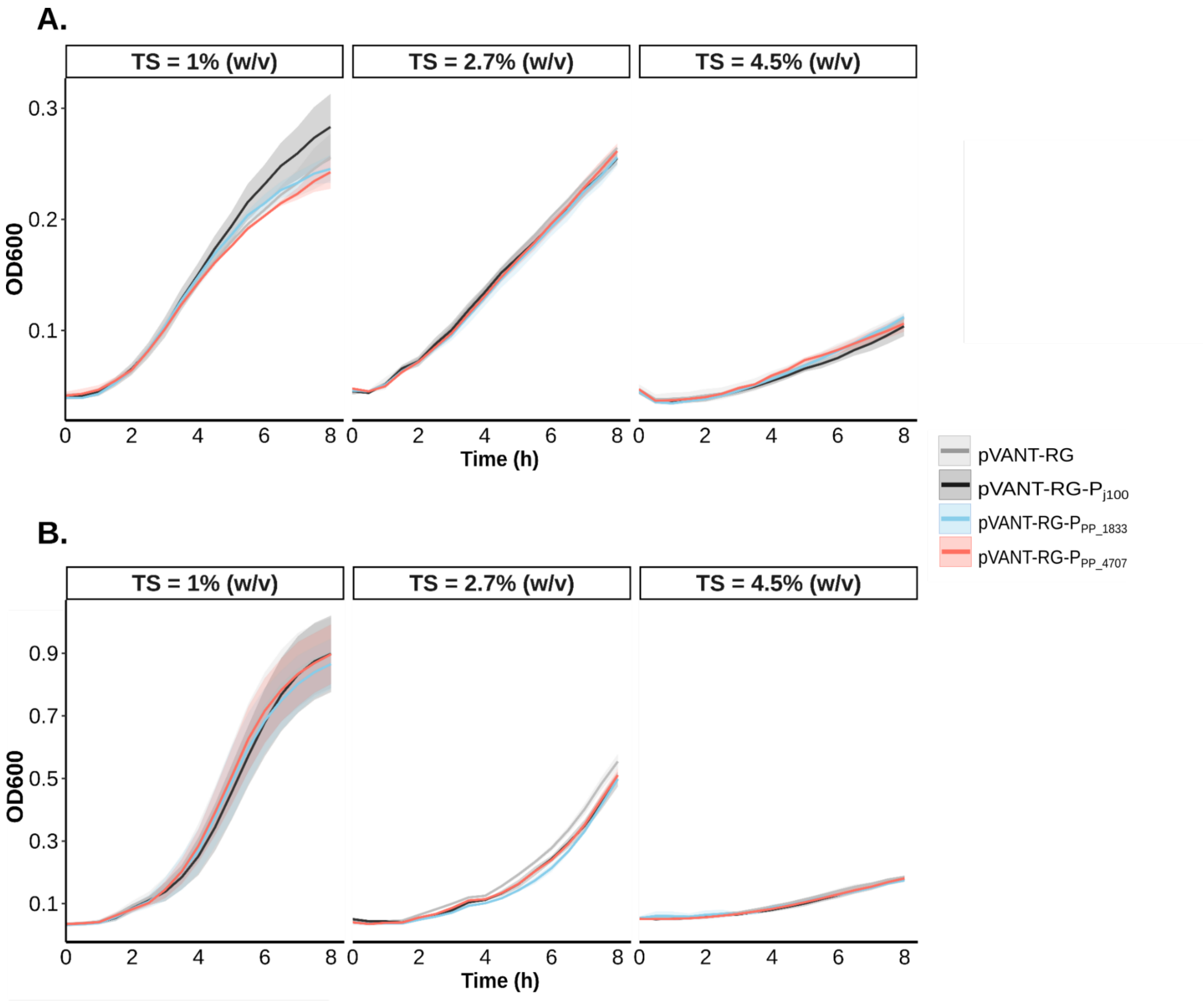


**Supplementary Figure 1. Evaluation of bacterial growth in different salt concentrations.** (A) Growth of *E. coli* DH10B and (B) *P. putida* KT2440. TS :Total salt concentration.

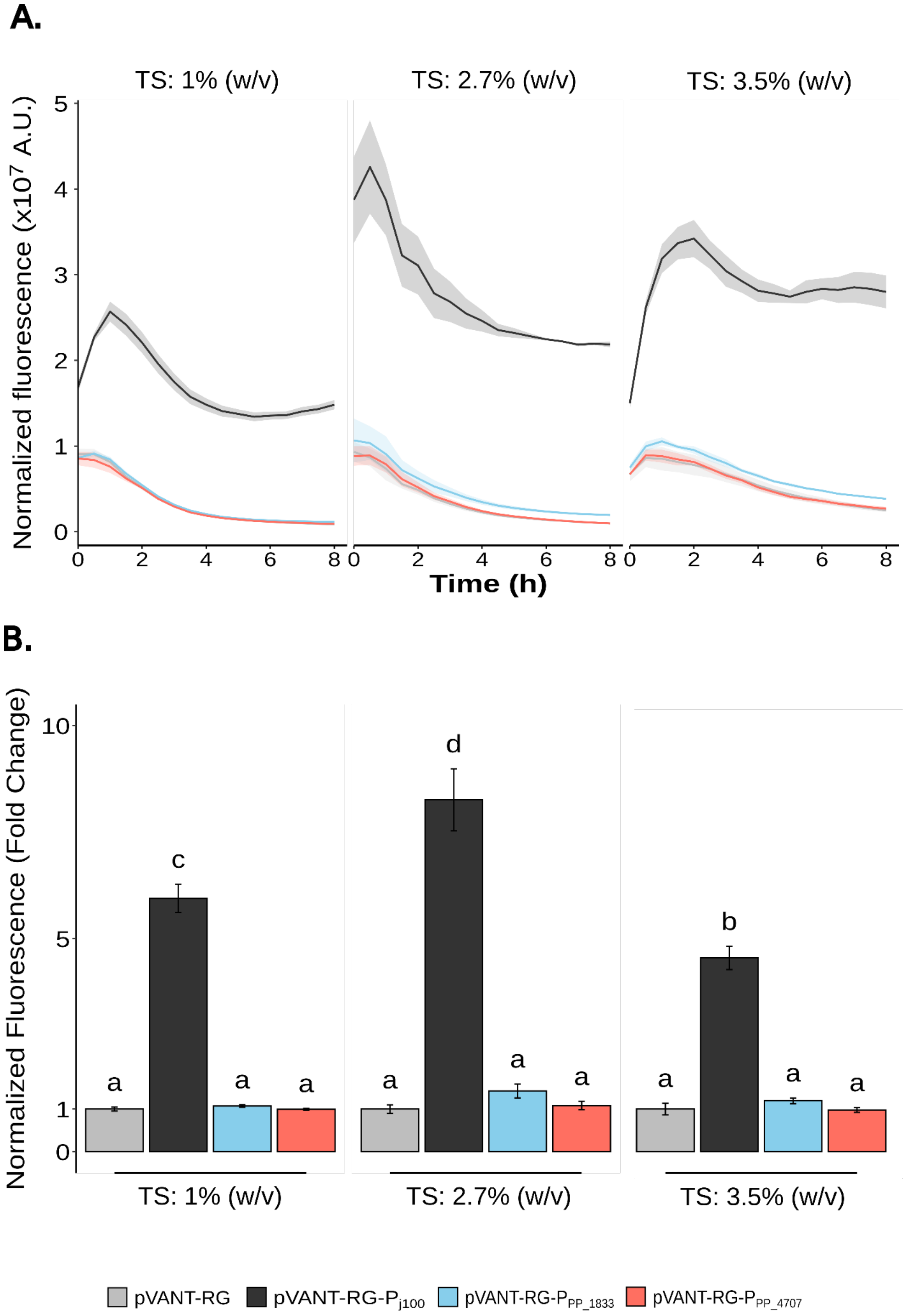


**Supplementary Figure 2. Variation of normalized fluorescence in *E. coli* DH10B.** Measurements were performed over 8 h (A) and at the third hour of incubation (B) in microplates containing minimal medium with glycerol and different NaCl concentrations. Two-way ANOVA followed by Tukey's post-hoc test revealed no statistically significant interaction between the plasmid lacking promoter regions (pVANT-RG) and the constructs containing the identified intergenic regions. A significant difference ($p < 0.05$) was observed only when samples were compared to the positive control, a strong promoter ($P_{j100}$) inserted upstream of the fluorescent protein (pVANT-RG-$P_{j100}$), as indicated by different letters above the bars. TS: 1% (w/v) = M9; TS: 2.7% (w/v) = M9 + 2.7% NaCl; TS: 4.5% (w/v) = M9 + 3.5% NaCl. A.U. = arbitrary units.

**Supplementary Table 1.** Primers used in this study for amplification of intergenic regions. Tm: Melting temperature.

| IGR | IGR Length (bp) | Primer Forward (PF) | PF Tm (°C) | Primer Reverse (PR) | PR Tm (°C) |
|---|---|---|---|---|---|
| PP_1833/PP_1834 | 266 | ATCAACTCCTGGCTGTTGTGTG | 55.18 | AGGGAACCTCCTGGCATAATTG | 54.08 |
| PP_4707/PP_4708 | 263 | GTGGGTCACTCCTGTTTTTCG | 53.65 | GCGAACGTCACGCAATAG | 51.69 |

**Supplementary Table 2.** Sequences of the IGRs investigated in this study based on the RefSeq NC_002947.4 genomic annotation

| IGR | Sequence |
|---|---|
| PP_1833/PP_1834 | ATCAACTCCTGGCTGTTGTGTGGCAGACATGAGTTGATGGTTGCAGTGACCGTGCCAGAATCCTGGCTGGCGAAAAACCCAGTAAATACAGGCGGTTACGGGGAATTTCCAGAAGCGGCATCGTGCAGGCTGCACGGTCGTGCATTTTGCACAGTGCATTTTGCACGGGTCAGGGTTTGCCTATCGGCTTCATAGAATCTGAACCTTTGACCGCAATGGCATGCCTGACGAACACTTGGGCCATCAATTATGCCAGGAGGTTCCCT |
| PP_4707/PP_4708 | GTGGGTCACTCCTGTTTTTCGAAAGGTCTGCGCCGTATGTCCTGGCGTCAGGTATGAAGGTAGTTGCACGGGCTGTGCCAGCTTTGCTGCTTCTGTTATTTCCTTTTAAATCAATAAGTTAGCAAATATCTATCACTTCGGTGGTCGTGCATTTTGCAATCAGCGCGGTAAACCTGCATGCAAGATGCAAGTCCGCAAAAGGTTCCCGTCCCCATGAAAAAAGGGCCCTGCGAAGGGCCCTTTCTCTATTGCGTGACGTTCGC |

## A.

AGGGAACCTCCTGGCATAATTGATGGCCCAAGTGTTCGTCAGGCATGCCATTGCGGTCAAAG
GTTCAGATTCTATGAAGCCGATAGGCAAACCCTGACCCGTGCAAAATGCACTGTGCAAAATG
CACGA**C**CGTGCAGCCTGCACGATGCCGCTTCTGGAAATTCCCCGTAACCGCCTGTATTTACT
GGGTTTTTCGCCAGCCAGGATTCTGGCACGGTCACTGCAACCATCAACTCATGTCTGCCACA
CAACAGCCAGGAGTTGATATG

## B.

GCGAACGTCACGCAATAGAGAAAGGGCCCTTCGCAGGGCCCTTTTTTCATGGGGACGGGAA
CCTTTTGCGGACTTGCATCTTGCATGCAGGTTTACCGCGCTGATTGCAAAATGCACGA**C**CACC
GAAGTGATAGATATTTGCTAACTTATTGATTTAAAAGGAAATAACAGAAGCAGCAAAGCTGGCA
CAGCCCGTGCAACTACCTTCATACCTGACGCCAGGACATACGGCGCAGACCTTTCGAAAAAC
AGGAGTGACCCACATG

**Supplementary Figure 3. Schematic representation of the promoters inserted in the intergenic regions.** Prediction of promoters contained in the intergenic regions: $P_{PP\_1833}$ (A) and $_{PP\_4707}$ (B). The regions of the -35 box (yellow), -10 box (green), Shine-Dalgarno sequence (blue), transcription start site (in bold), translation start site (pink), and promoter sequence (underlined) are shown. The translation start site (pink) lies outside the intergenic region, and both sequences are shown as reverse complements relative to the genome annotation.
